# Enzyme kinetics of CRISPR molecular diagnostics

**DOI:** 10.1101/2021.02.03.429513

**Authors:** Ashwin Ramachandran, Juan G. Santiago

## Abstract

CRISPR diagnostic assays have gained significant interest in the last few years. This interest has grown rapidly during the current COVID-19 pandemic where CRISPR diagnostics have been frontline contenders for rapid testing solutions. This surge in CRISPR diagnostics research prompts the following question: What exactly are the achievable limits of detection and associated assay times enabled by the kinetics of Cas12 and Cas13 enzymes? To address this question, we here present a model based on Michaelis-Menten enzyme kinetics theory applied to Cas enzymes. We use the model to develop analytical solutions for reaction kinetics and develop back-of-the-envelope criteria to validate and check for consistency in reported enzyme kinetics parameters. We applied our analyses to all studies known to us which report Michaelis-Menten-type kinetics data for CRISPR associated enzymes. These studies include all subtypes of Cas12 and Cas13 and orthologs. We found all studies but one clearly violate at least two of our three rules of consistency. We further use our model to explore ranges of reaction time scales and degree of reaction completion for practically relevant target concentrations applicable to CRISPR-diagnostic assays.

## 1. Introduction

There have been many interesting advancements in the field of CRISPR diagnostic assays in the last few years. This is particularly true during the current COVID-19 pandemic where CRISPR diagnostics have been frontline contenders for rapid testing solutions^1–4^. CRISPR diagnostic assays use subtypes of Cas12 or Cas13 enzymes for DNA or RNA detection, respectively^5–8^. The emerging and strong interest in CRISPR diagnostics research prompts the following question: What exactly are the achievable limits of detection (LODs) and associated assay times enabled by the kinetics of Cas12 and Cas13 enzymes (**Figure 1**)? As expected, the answer to this important question depends strongly on the CRISPR-associated enzyme kinetics as these establish and fundamentally limit the sensitivity of these diagnostics.

**Figure 1.**
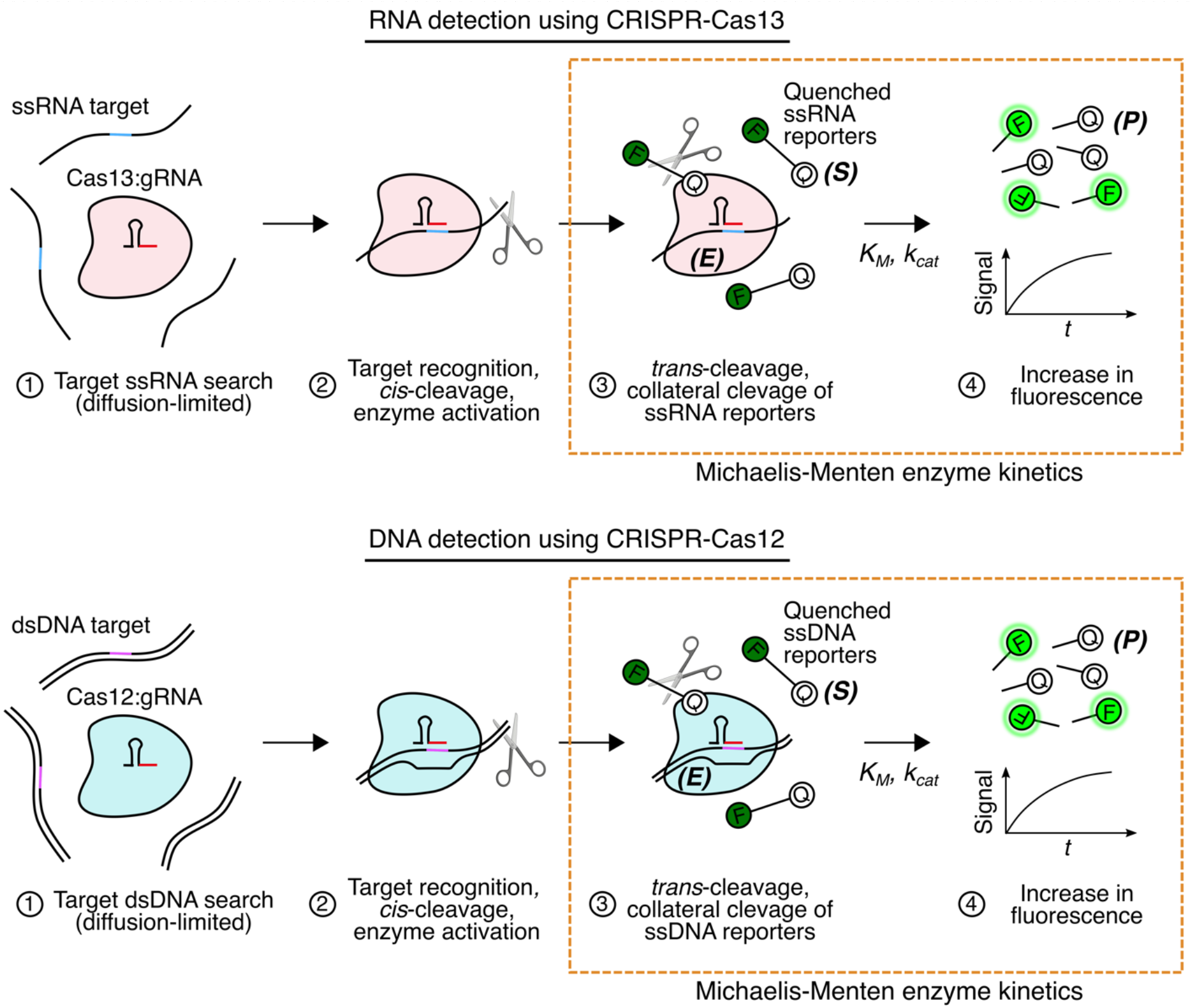
Schematic for RNA and DNA detection using CRISPR-Cas13 and Cas12, respectively. The Cas13:gRNA complex is activated by a target ssRNA (*cis*-cleavage; single turnover), after which the activated enzyme cleaves ssRNA molecules including ssRNA reporters indiscriminately (*trans*-cleavage; multi turnover). Cas12:gRNA can be activated by both target ssDNA and dsDNA (*cis*-cleavage; here, shown only for dsDNA). After activation, the Cas12 enzyme cleaves ssDNA molecules including ssDNA reporter molecules indiscriminately (*trans*-cleavage activity; multi turnover). The *trans*-cleavage activity for Cas12 and Cas13 each follows Michaelis-Menten kinetics, where the target-activated Cas-gRNA complex is the enzyme (*E*), the fluorescent reporter molecule is the substrate (*S*), and the cleaved reporter molecule is the product (*P*).

For ssDNA and dsDNA detection using CRISPR-Cas12, LbCas12a has been the most widely used Cas enzyme.^2,4,7,9–11^The study of Chen et al.^11^ was the first to characterize and report the kinetic rates of LbCas12a. The latter study reported an enzyme turnover rate (*k_cat_*) and Michalis-Menten constant (*K_M_*) of 250 per second and 4.94 × 10^−7^ M, respectively, with a catalytic efficiency (*k_cat_/K_M_*) of 5.1 × 10^8^ M^−1^s^−1^ for LbCas12a activated by target ssDNA. For a dsDNA activator, the study reported a *k_cat_* and *K_M_* of 1250 per second and 7.25 × 10^−7^ M, respectively, and a *k_cat_/K_M_* of 1.7 × 10^9^ M^−1^s^−1^. The latter value reportedly approached the limit of diffusion.^11^ Cofsky et al.^12^ studied the trans-cleavage activity of AsCas12a and reporter a *k_cat_* and *K_M_* of 0.6 per second and 2.7 × 10^−6^ M, respectively. We note the latter value of *k_cat_* is a factor 340 times lower than the second-lowest *k_cat_* ever reported for any CRIPSR associated enzyme (a study on LbuCas13a; **Table 1**).^13^

**Table 1.**
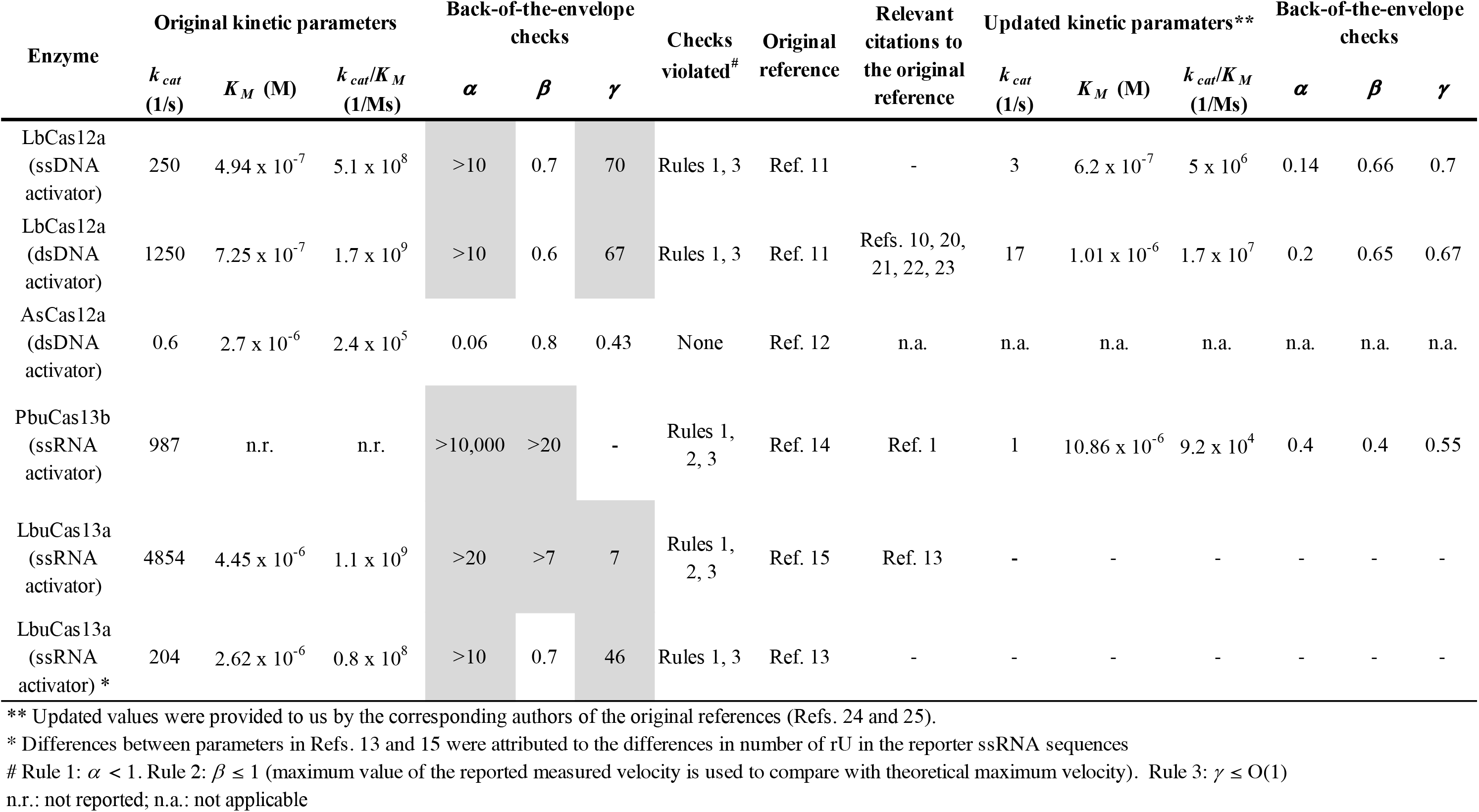
Summary of published enzyme kinetics parameters and our back-of-the-envelope checks for CRISPR-Cas12 and CRISPR-Cas13. Shaded boxes indicate violation of our back-of-the-envelope criteria for consistent kinetics data (Section 2.2). All but one study fail two or more validation rules.

| Enzyme | Original kinetic parameters |  |  | Back-of-the-envelope checks |  |  | Checks violated<sup>#</sup> | Original reference | Relevant citations to the original reference | Updated kinetic parameters** |  |  | Back-of-the-envelope checks |  |  |
| --- | --- | --- | --- | --- | --- | --- | --- | --- | --- | --- | --- | --- | --- | --- | --- |
| | $k_{cat}$ (1/s) | $K_M$ (M) | $k_{cat}/K_M$ (1/Ms) | $\alpha$ | $\beta$ | $\gamma$ | | | | $k_{cat}$ (1/s) | $K_M$ (M) | $k_{cat}/K_M$ (1/Ms) | $\alpha$ | $\beta$ | $\gamma$ |
| LbCas12a (ssDNA activator) | 250 | $4.94 \times 10^{-7}$ | $5.1 \times 10^8$ | >10 | 0.7 | 70 | Rules 1, 3 | Ref. 11 | - | 3 | $6.2 \times 10^{-7}$ | $5 \times 10^6$ | 0.14 | 0.66 | 0.7 |
| LbCas12a (dsDNA activator) | 1250 | $7.25 \times 10^{-7}$ | $1.7 \times 10^9$ | >10 | 0.6 | 67 | Rules 1, 3 | Ref. 11 | Refs. 10, 20, 21, 22, 23 | 17 | $1.01 \times 10^{-6}$ | $1.7 \times 10^7$ | 0.2 | 0.65 | 0.67 |
| AsCas12a (dsDNA activator) | 0.6 | $2.7 \times 10^{-6}$ | $2.4 \times 10^5$ | 0.06 | 0.8 | 0.43 | None | Ref. 12 | n.a. | n.a. | n.a. | n.a. | n.a. | n.a. | n.a. |
| PbuCas13b (ssRNA activator) | 987 | n.r. | n.r. | >10,000 | >20 | - | Rules 1, 2, 3 | Ref. 14 | Ref. 1 | 1 | $10.86 \times 10^{-6}$ | $9.2 \times 10^4$ | 0.4 | 0.4 | 0.55 |
| LbuCas13a (ssRNA activator) | 4854 | $4.45 \times 10^{-6}$ | $1.1 \times 10^9$ | >20 | >7 | 7 | Rules 1, 2, 3 | Ref. 15 | Ref. 13 | - | - | - | - | - | - |
| LbuCas13a (ssRNA activator) * | 204 | $2.62 \times 10^{-6}$ | $0.8 \times 10^8$ | >10 | 0.7 | 46 | Rules 1, 3 | Ref. 13 | - | - | - | - | - | - | - |

\*\* Updated values were provided to us by the corresponding authors of the original references (Refs. 24 and 25).
\* Differences between parameters in Refs. 13 and 15 were attributed to the differences in number of rU in the reporter ssRNA sequences

For ssRNA detection using CRISPR-Cas13, the study of Slaymaker et al.^14^ was the first to report enzyme turnover rates among all the subtypes and orthologs of Cas13. They reported a *k_cat_* of 987 turnovers per second for PbuCas13b. Subsequently, Shan et al.^15^ reported a *k_cat_* of 4854 per second and *K_M_* of 4.45 × 10^−6^ M using a 5 nt (of rU) ssRNA fluorescent reporter in an application of LbuCas13a for MicroRNA (miRNA) detection. Later, Zhou et al.^13^ showed for LbuCas13a that a shorter reporter molecule which was composed of 2 nt (of rU) lowered the *k_cat_* to 204 per second and *K_M_* of 2.62 × 10^−6^ M. Most recently, Fozouni et al.^1^ presented a study of SARS-CoV-2 viral RNA detection using LbuCas13a and a set of 3 gRNAs, and used a model to fit their experimental data which used parameters of *k_cat_* = 600 per second and *K_M_* between 1 × 10^−6^ M and 3 × 10^−6^ M.

In this work, we provide back-of-the-envelope analyses and scaling arguments based on Michaelis-Menten-type enzyme kinetics theory applied to CRISPR-diagnostics. We present a set of simple rules based on fundamental principles of species conservation, maximum reaction velocities, and reaction time scales, and these rules can be used as a check on reported parameters of enzyme kinetics. We apply these rules and perform calculations on several published studies. Surprisingly, we find all but one study (the aforementioned lowest ever reported value of *k_cat_*) clearly violate at least two of our three rules of consistency. Lastly, we develop and present results from analytical and numerical models for predicting Cas enzyme kinetics and use these to discuss the effect of kinetic parameters on the assay time scales and limits of sensitivity of CRISPR-diagnostic assays.

## 2. Theory

We first present an enzyme kinetics model for the trans-cleavage activity of CRISPR-associated enzymes. The mechanisms of relevant reactions are summarized in **Figure 1.** This model invokes Michaelis-Menten enzyme kinetics theory^16^ and can be used to develop closed-form approximations for product formation versus time. We then present three rules based on kinetics theory and fundamental principles of species conservations. These rules enable simple back-of-the-envelope calculations and can be performed to check the self-consistency and/or accuracy of reported enzyme kinetics data.

### 2.1 Michaelis-Menten model for *trans*-cleavage activity

Upon recognition of the target nucleic acid, the CRISPR-enzymes are activated and exhibit non-specific trans-cleavage of single-stranded nucleic acids (c.f. **Fig. 1**). ^9,11,17^ This trans-cleavage activity is a multi-turnover enzymatic reaction catalyzed by the target activated CRISPR enzyme. This reaction is the key process underlying nucleic acid detection assays. Cas12 can be activated by target ssDNA and dsDNA, while Cas13 is activated by ssRNA. Upon activation, Cas12 and Cas13 indiscriminately cleave ssDNA and ssRNA molecules, respectively. CRISPR assays typically use fluorescent nucleic acid molecules (with a quencher) as reporters, which fluoresce only when they are cleaved by the CRISPR enzyme in the presence of target.

The trans-cleavage CRISPR enzymatic reaction can be modeled as

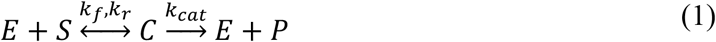

where, *E* is target-activated Cas12/13-gRNA complex, *S* is the uncleaved reporter, *C* is the reaction intermediate (enzyme-substrate reporter complex), *P* is the cleaved reporter molecules, *k_f_* and *k_r_* are the forward and reverse rate constant for the reaction between *E* and *S,* and *k_cat_* is the catalytic turnover rate of the enzyme. The reaction rate laws for the individual species are as follows:

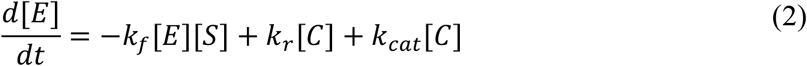

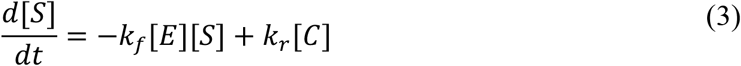

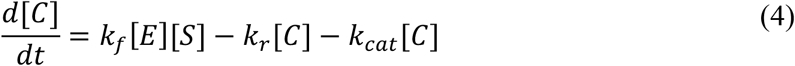

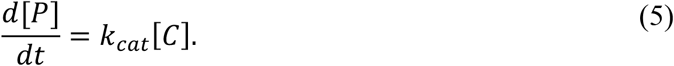

These are subject to initial conditions *[E]*(0) = *E*_0_, [*S*](0) = *S*_0_, [*C*](0) = 0, and [*P*](0) = 0. Next, we use the quasi-steady state assumption (see Briggs-Haldane approximation^16^) which models the concentration of the intermediate enzyme-substrate complex as nearly constant in time. This powerful simplification enables us to write

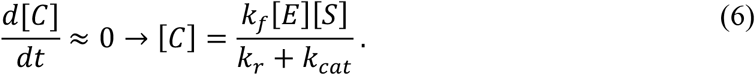

Further, species conservation yields

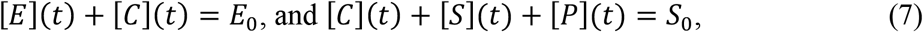

where *E*_0_ and *S*_0_ are respectively the initial amount of target-activated enzyme and substrate (most typically a synthetic nucleic acid reporter molecule). The “*t*” in parentheses indicate time-dependent quantities. We note that *E*_0_ is at most equal to the amount of target nucleic acid present in the sample (usually the Cas-gRNA complex is present in abundance). However, at very low concentrations of target, target recognition and activation of the CRISPR enzyme (*cis*-cleavage) is limited by the diffusion of the target molecules to the active site of the enzyme.^18^ For simplicity, we will here assume that *E*_0_ is given by the concentration of the target nucleic acid.

Under these assumptions, the rate of product formation (reporter cleavage) is governed by the Michaelis-Menten equation

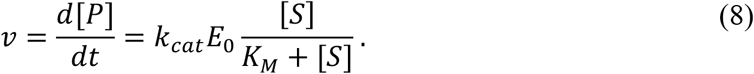

In traditional Michaelis-Menten kinetics systems, *E*_0_ ≪ *S*_0_ and the kinetics are studied over short time scales (relative to the overall product formation time scale). Hence, in this limit, very little product is formed *P* ≪ *S*_0_ and [*S*] ≈ *S*_0_. Thus, the product is formed at a constant rate (e.g., linear fluorescence increase vs time), and the reaction velocity is given by Equation (8) with [*S*] ≈ *S*_0_

Further, in most practical applications of CRISPR-diagnostics, the concentration of the substrate is much smaller than the Michaelis-Menten constant of the enzyme, [*S*] ≪ *K_M_*. Typically 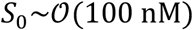, and 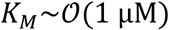. Thus, we can use this approximation to further derive an expression for the evolution of the product (reporter cleavage) versus time, over much longer time scales. Specifically, we can rewrite Equation (8) as

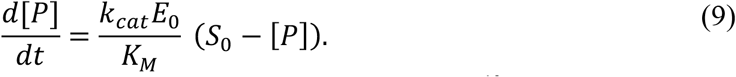

Equation (9) can be rearranged into a linear first-order differential equation^19^ as

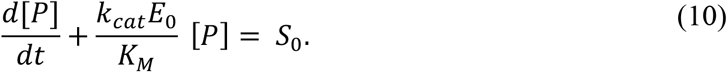

Equation (10) can be solved analytically for the concentration of product formed (reporters cleaves) versus time as

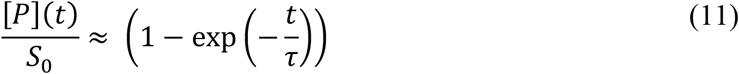

where *τ* = *K_M_/k_cat_E*_0_. Here *τ* is a time scale that governs the characteristic time to approximately complete the reaction, and [*P*](*t*)/*S*_0_ is simply the fraction of cleaved reporters relative to the total initial uncleaved reporters. Physically, we see that the time scale of significant product formation *τ* decreases with: (i) higher turnover rate *k_cat_*, (ii) higher target concentration *E*_0_, (iii) lower *K_M_* (which represents a higher affinity of the activated enzyme to the substrate). Further, we also see that the ratio of *k_cat_* to *K_M_* is very important for the product formation rate and this ratio arises naturally from the current analysis.

### 2.2 Back-of-the-envelope checks on reported kinetics data

We here present three simple back-of-the-envelope hand calculations that can be used to verify the accuracy of reported kinetics data from experiments and/or simulations. The first rule derived here originates simply from species conservation: The amount of product formed during the initial linear portion *t_lin_* of the kinetics measurements (c.f. **Figure 2**; early time scales where *τ* < 1) cannot exceed the initial amount of substrate (before the start of the reaction). *t_lin_* is a quantity that typically is easily apparent from experimental data (e.g. the duration of the initial, nearly linear increase of fluorescence versus time data). We can define the non-dimensional parameter *α* as the ratio of the total product formed at *t_lin_* and the initial substrate concentration *S*_0_. For most CRISPR diagnostics assays, *α* is the ratio of the amount of cleaved reporters at time *t_lin_* to the initial concentration of the uncleaved reporters *S*_0_. Simple conservation of species then limits this value to unity as in

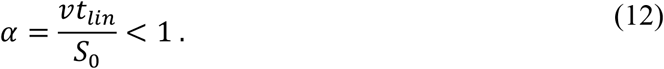

**Figure 2.**
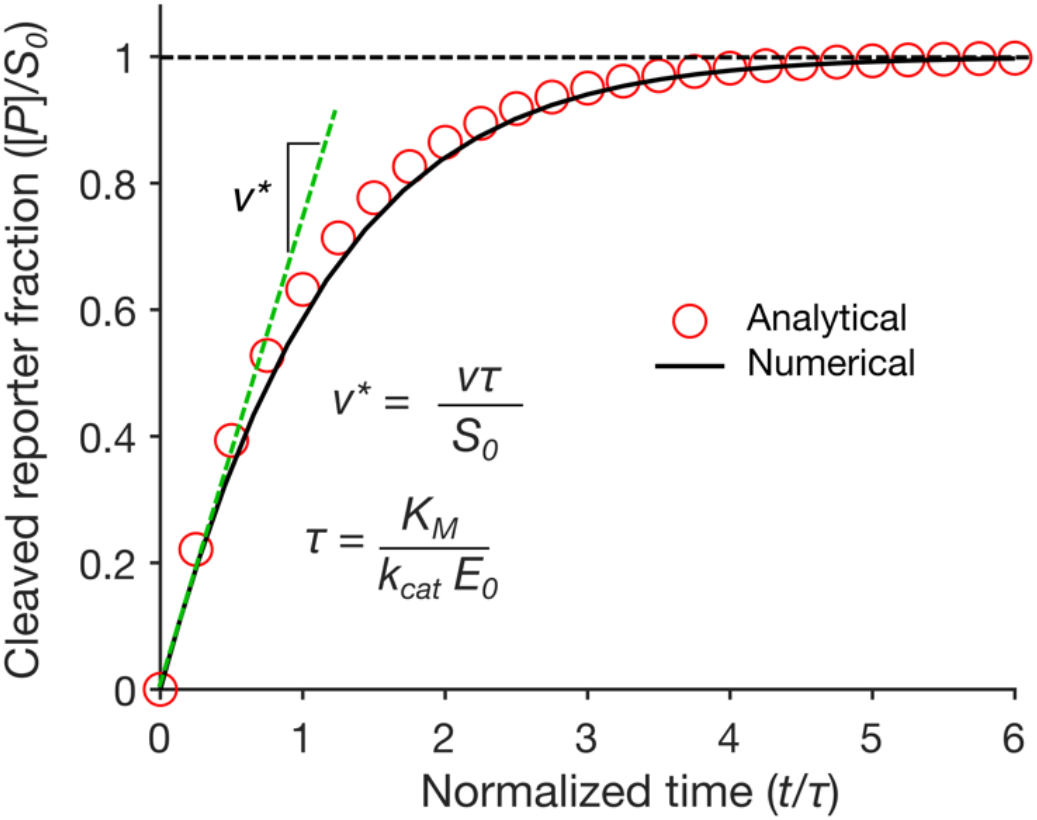
Fraction of reporters cleaved by the Cas enzyme versus time normalized by the reaction time scale, *τ*, as predicted by the numerical model (solid line) and analytical solutions (symbols). Results are shown for kinetic parameters *k_cat_* and *K_M_* of 17 s^−1^ and 1.01 × 10^−6^ M, and initial target­ activated enzyme concentration of 100 pM and initial uncleaved reporter concentration of 200 nM.

Here *v* is the reported reaction velocity (e.g., in nM/s).

The second rule is derived from the theoretical maximum reaction velocity possible as given by Michaelis-Menten kinetics. From Equation (8), the maximum rate of product formation occurs when the substrate concentration is far in excess compared to *K_M_*, [*S*] ≫ *K_M_*, and the theoretical maximum velocity *v_max_* is equal to *k_cat_E*_0_. One can therefore define a non-dimensional quantity, *β* as the reported velocity *v* normalized by *v_max_* as follows:

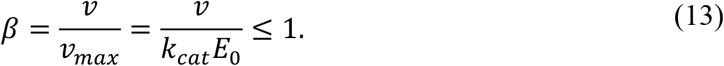

Third, we observe that the time scale of linear portion of the kinetics data *t_lin_* should be at most on the same order as the reaction time scaler. For this, we define a non-dimensional quantity, *γ*, defined as the ratio of *t_lin_* and *τ*, and so the third rule is given by

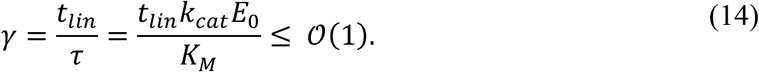

Lastly, we note that the first rule (condition on *α*) does not require *a priori* estimates of the reaction parameters *k_cat_* and *K_M_*. The first rule is therefore a good check on ensuring consistent fluorescence versus concentration calibration for experiments. For example, inconsistencies in calibration or conversion from fluorescence to concentration will result in unphysical values of *v*, and these artifacts can lead to an apparent violation of the first rule (hence, we here assume the initial substrate concentrations and times are properly reported). The second and third rules rely on empirically determined (e.g. by fitting models to experimental data) reaction parameters *k_cat_* and *K_M_* (e.g., using a curve fit to initial reaction velocity versus substrate concentration kinetics data). The latter two rules (involving *β* and *γ* provide useful validations of self-consistency in kinetics data and the reported values of *k_cat_* and *K_M_*. Briefly, all data should obey Rules 1 and 2, and reactions for which [*S*] ≪ *K_M_* (this inequality can be interpreted as less than about 5 fold) must obey Rule 3.

## 3. Results and Discussion

### 3.1 Back-of-the-envelope checks on published data

Presented in **Table 1** are the back-of-the-envelope checks for quantities *α, β*, and *γ* (c.f. Section 2.2) which we performed on published studies that present Michaelis-Menten-type characterization experiments and kinetics data for different Cas12 and Cas13 enzymes. To the best of our knowledge, we have included all published studies to date which present any Michaelis-Menten kinetics data for CRISPR-Cas 12/13. We found that all of the studies except one listed in the first column of **Table 1** violated at least two of the three checks involving *α, β*, and *γ* (see fifth, sixth, and seventh columns, respectively). For example, the studies of Chen et al.^11^ for LbCas12a and Zhou et al.^13^ for LbuCas13a both had values of *α* and *γ* significantly greater than 1. The study of Shan et al.^15^ for LbuCas13a reported values of *α, β*, and *γ* which were each greater than 1, and so violated all the three rules. Slaymaker et al.^14^ studied PbuCas13b, and the study violated at least two (*α* and *β* conditions) out of the three checks. The study of Cofsky et al.^12^ involving AsCas12a was the only one which passed all three checks. Interestingly, the study of Cofsky et al.^12^ is a dramatic outlier in the reported value of *k_cat_* (c.f. second column of **Table 1**). The results in **Table 1** suggest that the kinetics parameters (second, third, and fourth columns of **Table 1)** published in the aforementioned studies (first column of **Table 1)** with at least one check violated are not reliable and the data may be grossly incorrect. Also listed in **Table 1** are other relevant publications^1,10,20–23^ (column 10) which cite the aforementioned studies and either directly rely (e.g. in making estimates or modeling experiments) on the published kinetics data or provide new experimental data which reportedly corroborates the older kinetics parameter values that violated our checks.

Lastly, we present updated kinetics parameter values (columns 11, 12 and 13 of **Table 1)** which we recently received from the corresponding authors of Refs.^11,14^ through personal communication ^24,25^. The current authors learned from the corresponding authors of Ref.^11^ (the oldest study in the set) that the error in kinetic rates of LbCas12a in their original publication was due to an incorrect conversion of relative fluorescence units to molar concentration for the initial reaction velocity calculations. Unlike most of originally published data, the updated kinetic rates for Refs.^11,14^ are consistent with (i.e. pass) our three back-of-the-envelope checks (see columns 14, 15, and 16 of **Table 1).** That is, for these updated values, the three quantities *α, β,* and *γ* are each less than unity as expected. If these updated kinetic parameters are correct, a major consequence is that the kinetics of LbCas12a is not catalytically efficient or diffusion-limited (*k_cat_/K_M_*~10^6^ to 10^7^ M^−1^ s^−1^), contrary to wide consensus in the CRISPR field (e.g. see Refs. ^4,10,11,20,22^). Also, turnover rates for LbCas12a and PbuCas13b, respectively, which are 2 to 3 orders of magnitude lower would imply that proportionally longer reaction time scales are required for *trans*-cleavage activities (Equation (11)). As we discuss in the next section, this would have dramatic effects on the achievable limits of detection using CRISPR diagnostics (e.g. without pre-amplification).

### 3.2 Predictions of analytical and numerical enzyme kinetics model

We here use the model developed in Section 2.1 to explore the kinetics of Cas enzymes. **Figure 2** shows the fraction of reporters cleaved by the target-activated enzyme divided by the initial amount of uncleaved reporters present in the reaction versus time normalized by the reaction time scale *τ*. Results are shown for *k_cat_* and *K_M_* of 17 s^−1^ and 1.01 × 10^−6^ M, which are based on the updated kinetic parameters for LbCas12a enzyme and a dsDNA activator **(Table 1).** Also, we assumed 100 pM (*E*_0_) of target dsDNA and 200 nM (*S*_0_) of ssDNA reporters for **Figure 2.** Presented in **Figure 2** are results from both a numerical model that solves the complete set of coupled differential equations for the enzyme kinetics (Equations (2)-(5)) (solid line) and the analytical solution (shown as symbols in **Figure 2)** given by Equation (11). First, we see very good agreement between the analytical and numerical solution results. Also, indicated by the green dashed line is a tangent to the numerical solution at early reaction times (*t* → 0 s), and the slope of this line is the scaled reaction velocity *v*\* (= *vτ/S*_0_), where *v* is a quantity typically measured in Michaelis-Menten-type kinetics analysis (Equation (8)). Consistent with our normalization and species conservation (*α* condition; **Section 2.2),** note that the cleaved reporter fraction asymptotes to unity for large times. Also, note from **Figure 2** that *τ* (= *K_M_/k_cat_E_o_*) represents the physically relevant product formation/reaction time scale. Consistent with our formulation (i.e. the *γ* condition; **Section 2.2),** we see that most of the early-time linear increase in the product amount happens for *t/τ* < 1. Lastly, we note that, due to our choice of normalization variables, the curve showed in **Figure 2** can be fairly generalized to other concentrations of target and reporters (see Equation (11)) as long as *E*_0_ ≪ *S*_0_, and for *S*_0_ lower than *K_M_* by at least a factor of ~5.

Next, we study the fraction of cleaved reporters as a function of the target nucleic acid concentration and the concentration of the initial uncleaved reporter **(Figure 3).** We assume the same enzyme kinetics parameters as **Figure 2,** and study the product formed after 60 min of reaction time (commensurate with typical molecular diagnostic assays). First, note that, for initial reporter concentrations lower than about 1 μM, the fraction of cleaved reporters is nearly invariant with the initial reporter concentration (this limit is provided by the analytical solution in Equation (11)). Further, the fraction of cleaved reporters after 60 min reaction varies significantly from ~100% at target concentrations greater than ~100 pM to less than 1% for target concentrations lower than ~200 fM. Lastly, we note that the cleaved reporter fraction shown in **Figure 3** can be interpreted as an estimate measure of the signal-to-noise ratio of assays that rely on end-point fluorescence readout for target detection (e.g. assuming a sensitive detector with a low noise). The reason for this is that uncleaved reporters very likely represent a substantial degree of background signal (e.g. due to imperfect quenching, probe degradation, and/or unwanted free dye).

**Figure 3.**
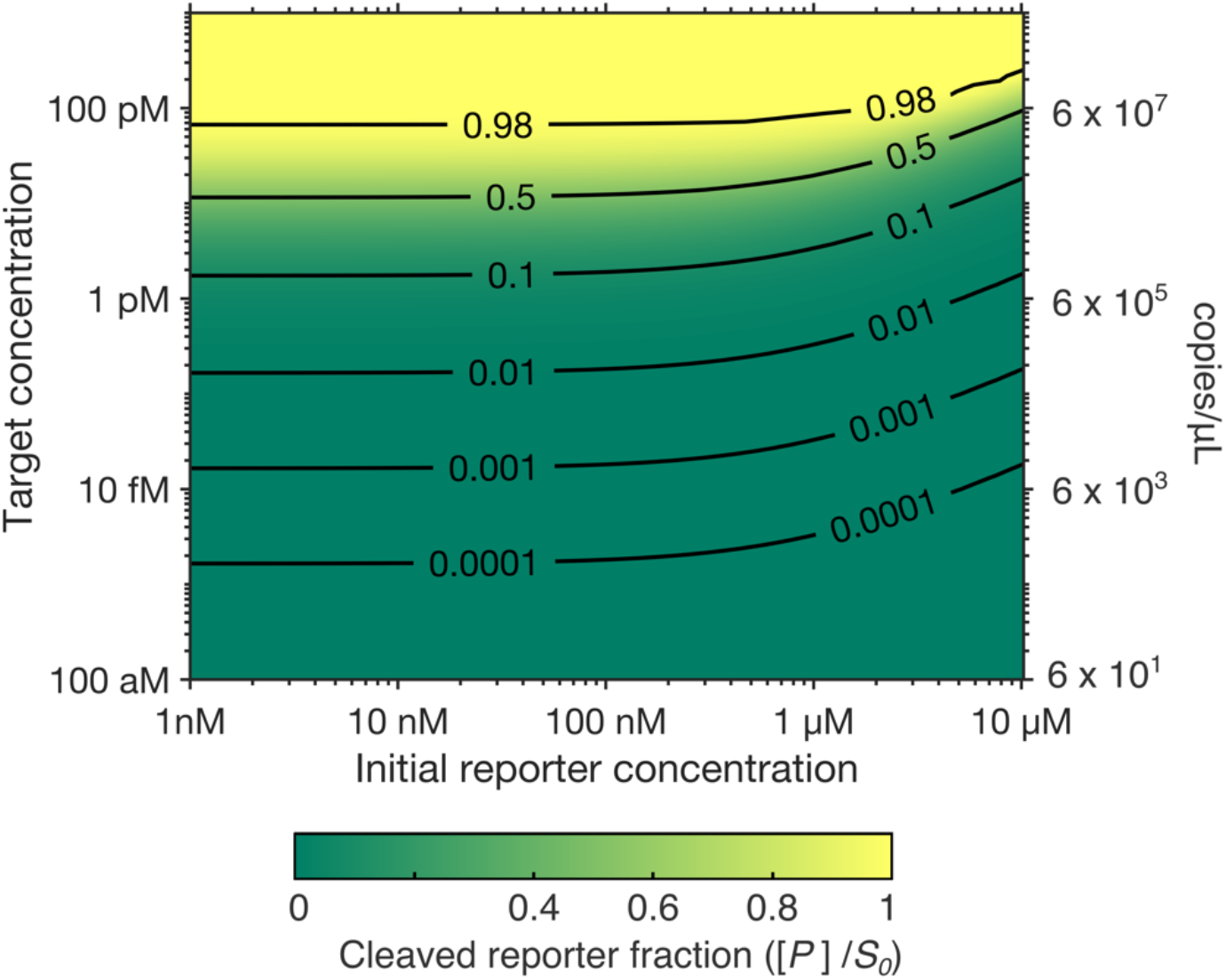
Model predictions for the fraction of cleaved reporters [*P*] relative to the initial amount of uncleaved reporters *S*_0_. Contours of the cleaved reporter fraction are plotted as a function of target nucleic acid concentration and initial uncleaved reporter concentration, for a 60 min assay time, and *k_cat_* and *K_M_* of 17 s^−1^ and 1.01 × 10^−6^ M, respectively (same as **Figure 2).** Contour levels of 0 and 1 represents no reporter cleavage and complete reporter cleavage, respectively.

### 3.3 Scaling of assay time scales

Here, we present the variation of reaction time scale *τ* for varying amounts of target nucleic acid based on Michaelis-Menten kinetics analysis (Section 2.1). **Figure 4** shows contours of *τ* (in seconds) as a function of the Cas enzyme catalytic efficiency (*k_cat_/K_M_*) and the target concentration *E*_0_. Note the inversely proportional relationship for the time scale between catalytic efficiency and target abundance: 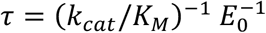. High catalytic efficiency and large target concentrations lead to shorter reaction times scales (hence faster reaction completion and fast detection). Notably, if the updated kinetics parameters in **Table 1** are true, the time scale for *trans*-cleavage reactions involving LbCas12a (for which *k_cat_/K_M_* ~ 10^6^ to 10^7^ M^−1^s^−1^) can vary from ~5 min for target concentrations greater an 1 nM to ~1 day for target concentrations less than ~1 pM. Note that, by definition, the reaction time scaler denotes the time at which about ~63% of the reporters are cleaved.

**Figure 4.**
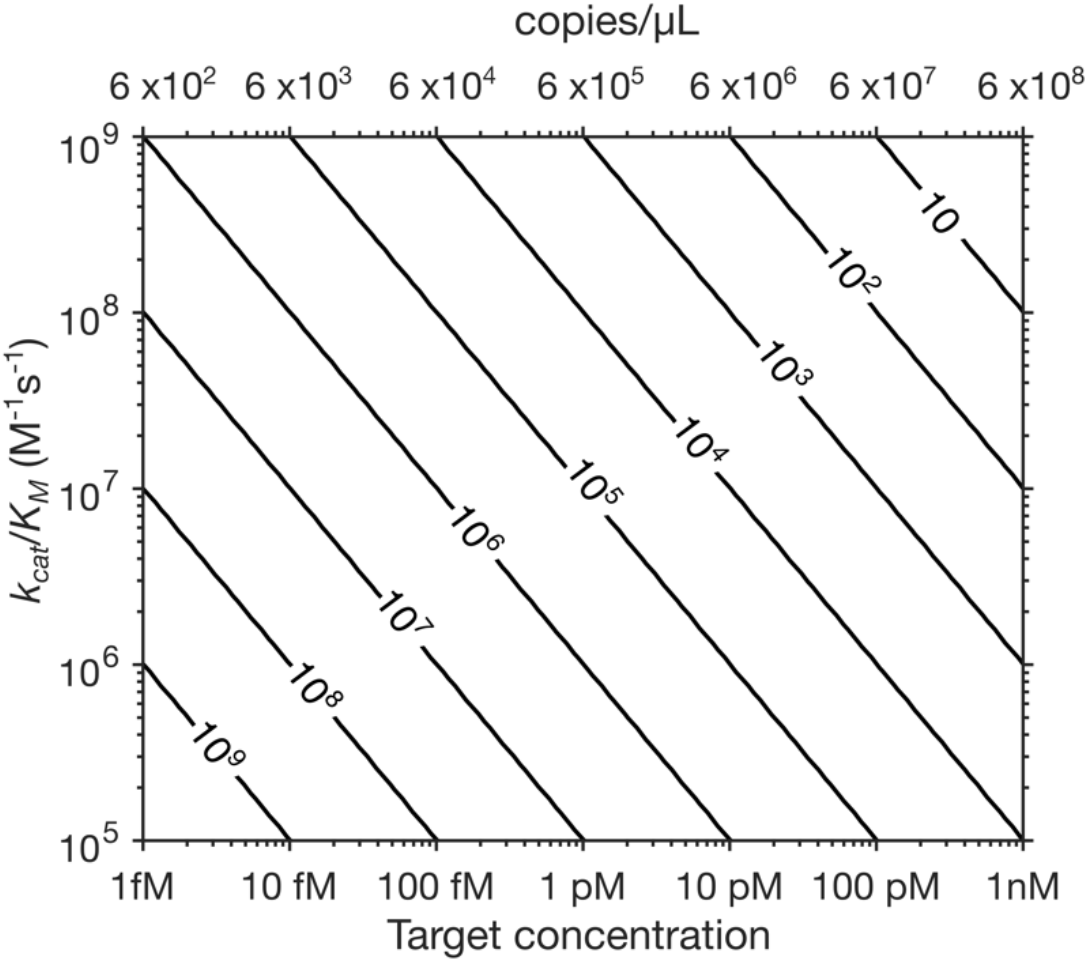
Contours of the reaction time scale *τ* (in seconds) versus the Cas enzyme catalytic efficiency parameter *k_cat_/K_M_* and the target nucleic acid concentration *E*_0_. Target concentrations of order 10,000 copies per μL or lower are likely very difficult to detect in a 60 min assay.

## 4. Summary

We presented a model for the trans-cleavage of CRISPR-associated enzyme kinetics based on Michaelis-Menten kinetics theory. The model is used to develop simple, back-of-the-envelope rules to check for consistency in experimentally derived and reported kinetic rate parameters. We applied these rules to data reported in published studies which characterize CRISPR-enzyme *trans*-cleavage rates. We found that all except one of the published studies which present Michaelis-Menten kinetics data violated these basic validation criteria. The single study which passed our three validation rules reported a *k_cat_* value which is a dramatic outlier relative to all other published values of *k_cat_*. We further used our model to develop simple scaling laws relating the kinetic parameters to reaction time scales and the degree of reaction completion. We applied this theory, using updated kinetic rates for LbCas12a, to make predictions of enzyme kinetics across practically relevant ranges of target concentrations for CRISPR-detection.

